# p53 restoration suppresses retrotransposon-driven chromosomal instability through nonlinear let-7 feedback and stochastic burst control

**DOI:** 10.64898/2026.02.27.708419

**Authors:** L. Boominathan

## Abstract

Long-read sequencing has revealed that concurrent LINE-1 (L1) retrotransposition events on non-homologous chromosomes frequently generate reciprocal chromosomal translocations early in tumorigenesis, establishing retrotransposons as active drivers of structural genome evolution (1). Endogenous mechanisms that constrain L1-mediated chromosomal instability remain incompletely defined. p53 transcriptionally induces tristetraprolin (TTP/ZFP36) and let-7 microRNAs, both directly and indirectly via repression of the MYC–LIN28 oncogenic axis (2). Mature let-7 suppresses human L1 retrotransposition by binding L1 mRNA and impairing ORF2p translation (3). Here we integrate these findings into a nonlinear dynamical systems model linking p53 activation, MYC–LIN28–let-7 feedback, and L1 RNA kinetics. Deterministic analysis uncovers bistability, with a sharp p53 activation threshold separating genome-unstable (high L1) and genome-stable (low L1) attractor states. Stochastic simulations reproduce the punctuated, clustered insertion patterns observed in tumors. Modest p53 restoration disproportionately collapses burst frequency, reducing cumulative structural rearrangement burden—including reciprocal translocations—by >70% under moderate assumptions. These results reposition p53 restoration as a threshold-dependent, retrotransposon-restrictive strategy to limit early genomic diversification and clonal evolution in cancer, with implications for pharmacologic reactivation therapies.

## Introduction

Nearly half the human genome derives from transposable elements, with LINE-1 (L1) as the sole autonomously active retrotransposon in humans. Although largely quiescent in most somatic cells through epigenetic silencing (DNA methylation, histone modifications, piRNAs, and KRAB-ZFPs), L1 elements can become derepressed under specific conditions such as early embryogenesis, germ-line development, or pathological states including cancer. Recent long-read sequencing studies have revealed that tumors exhibiting high L1 activity harbor thousands of somatic insertions, the majority of which are 5′-truncated and non-autonomous. Critically, concurrent retrotransposition events occurring on non-homologous chromosomes frequently generate reciprocal chromosomal translocations—balanced structural rearrangements—at rates of approximately one rearrangement per 40–60 somatic insertions in cases with hyperactive L1 landscapes (1). These events often precede whole-genome doubling and contribute to early chromosomal instability, positioning retrotransposition as a structural driver that operates independently of—or in concert with—replication stress or telomere crisis (7).

### LINE-1 Retrotransposition Mechanisms

Full-length, retrotransposition-competent human L1 elements (primarily from the L1Hs subfamily) are ∼6 kb in length and consist of a 5′ untranslated region (UTR) containing an internal RNA polymerase II promoter, two open reading frames (ORFs), and a 3′ UTR with a poly(A) tail. ORF1 encodes ORF1p, a ∼40 kDa homotrimeric RNA-binding protein with nucleic acid chaperone activity that facilitates RNP formation and RNA unwinding. ORF2 encodes ORF2p, a ∼150 kDa multifunctional protein possessing an endonuclease (EN) domain and a reverse transcriptase (RT) domain essential for mobilization. L1 retrotransposition proceeds via a target-primed reverse transcription (TPRT) mechanism:

1. Transcription: Derepressed L1 elements are transcribed into full-length, polyadenylated bicistronic mRNA.
2. Translation and RNP Assembly: The mRNA is exported to the cytoplasm, where ORF1p and ORF2p are translated. ORF1p trimers bind the L1 RNA with high affinity (often exhibiting cis-preference), and together with ORF2p form ribonucleoprotein (RNP) particles that may undergo liquid-liquid phase separation.
3. Nuclear Import: The RNP translocates into the nucleus.
4. Target Site Nicking: ORF2p’s EN domain introduces a single-strand nick in the bottom DNA strand at relaxed consensus sequences (typically 5′-TTTT/AA-3′ or variants in AT-rich regions), exposing a free 3′-OH group that base-pairs with the L1 mRNA poly(A) tail.
5. Target-Primed Reverse Transcription (TPRT): ORF2p’s RT domain extends from the 3′-OH using the L1 RNA as template, synthesizing the first-strand cDNA directly at the genomic site. Synthesis is often abortive, resulting in 5′-truncated insertions (∼90–95% of somatic events are <1 kb). 7
6. Second-Strand Cleavage and Integration: ORF2p nicks the complementary strand; host DNA repair pathways (e.g., non-homologous end joining) complete second-strand synthesis and ligation, generating characteristic target site duplications (TSDs) of 7–20 bp flanking the new insertion.

Most insertions occur in gene-poor, AT-rich genomic regions (10), though non-canonical, EN-independent pathways can exploit existing DNA breaks, replication forks, or induced damage. In tumors with elevated L1 activity, the propensity for concurrent (often synchronous) events on non-homologous chromosomes directly promotes reciprocal translocations, reshuffling large genomic segments and contributing to oncogene activation, tumor suppressor inactivation, and accelerated clonal evolution (1).The tumor suppressor p53 orchestrates classical responses to cellular stress, including cell cycle arrest, apoptosis, senescence, and DNA repair. Beyond these functions, p53 exerts post-transcriptional control through microRNAs and RNA-binding proteins. p53 transcriptionally induces tristetraprolin (TTP/ZFP36), an AU-rich element-binding protein that destabilizes oncogenic transcripts such as MYC and LIN28A/B, as well as members of the let-7 microRNA family, both directly (via promoter binding) and indirectly by disrupting the MYC–LIN28 oncogenic axis (2). MYC transactivates LIN28, while LIN28 inhibits let-7 maturation by binding pri-/pre-let-7 and promoting oligouridylation and decay, creating a mutual antagonism loop. p53 activation breaks this loop: miR-34a and miR-145 (induced by p53) repress MYC, TTP destabilizes MYC and LIN28 mRNAs, and direct p53 induction of let-7 further amplifies mature let-7 levels.Mature let-7 suppresses L1 retrotransposition by binding conserved sites in L1 mRNA, recruiting Argonaute-containing RISC complexes to impair translation of ORF2p without significantly affecting L1 transcription or ORF1p levels (3) (figure 1). Reduced ORF2p limits RNP assembly, reverse transcription efficiency, and new insertion events, thereby constraining the burst dynamics that drive concurrent translocations.We sought to integrate these molecular findings into a nonlinear dynamical systems framework to test whether p53 activation imposes threshold-dependent, bistable suppression of L1 bursts and associated structural rearrangements, including reciprocal translocations. By modeling the p53–TTP/miR-34a/145– MYC–LIN28–let-7–L1 RNA network, we examine whether modest restoration of p53 activity can disproportionately stabilize the genome through collapse of retrotransposition frequency and burst amplitude.

**Figure 1.**
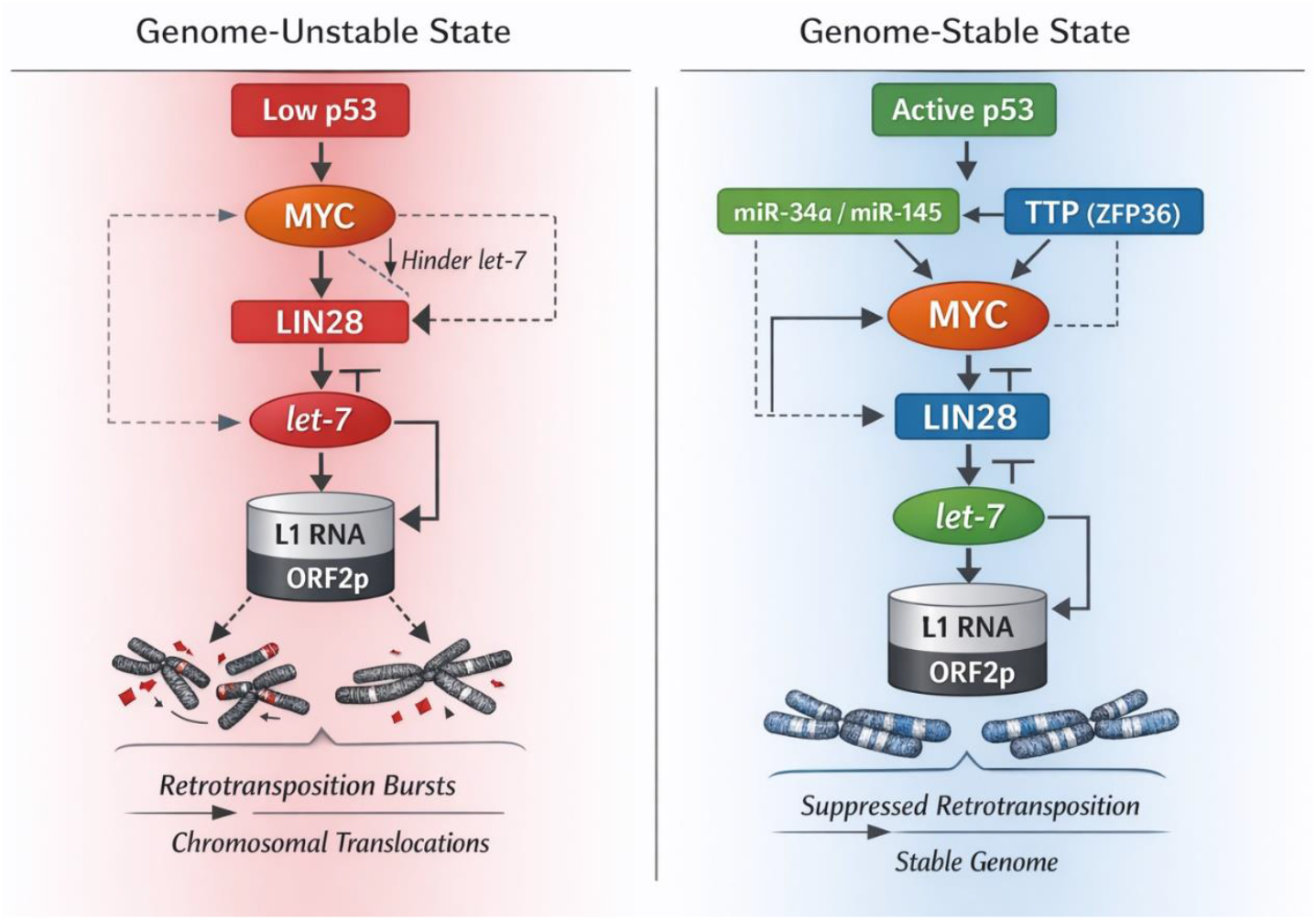
Mechanistic network diagram. **p53 activation establishes a bistable switch suppressing LINE-1 retrotransposition via let-7 feedback. Schematic representation of the core regulatory network in two contrasting states. Left panel (genome-unstable state): Low p53 activity sustains high MYC (via absent miR-145 repression) and high LIN28 (via TTP deficiency), resulting in low mature let-7 levels, elevated L1 mRNA and ORF2p, and frequent concurrent retrotransposition bursts leading to reciprocal chromosomal translocations. Right panel (genome-stable state): p53 activation induces TTP/ZFP36 and miR-34a/145, repressing MYC (direct miR-145 targeting of 3′ UTR plus TTP-mediated mRNA destabilization) and LIN28 (TTP destabilizes LIN28A/B transcripts), thereby relieving let-7 maturation block and enabling direct p53 transcriptional induction of let-7 family members; elevated let-7 binds L1 mRNA, impairing ORF2p translation and suppressing bursts. Solid arrows denote activation or induction; T-bars indicate repression or inhibition; dashed lines highlight feedback loops (MYC → LIN28 ┤ let-7 ┤ MYC double-negative motif); red highlights denote p53 inputs and let-7 effector arm; gray shading emphasizes L1 RNA/ORF2p levels. Network topology integrates established mechanisms.**

## Methods

### ODE system

The reduced deterministic model describes the core dynamics of the p53-regulated network suppressing LINE-1 (L1) retrotransposition. Species are normalized to dimensionless quantities (0–1 scale for relative concentrations/activities): oncogenic axis O (composite MYC–LIN28 levels), let-7 microRNA L, L1 RNA R. p53 activation U serves as the bifurcation control parameter (normalized 0 = complete loss, 1 = full wild-type restoration). Full equations incorporate Hill functions to capture cooperative nonlinearities in repression, maturation relief, and let-7-mediated decay:dO/dt = k_O / (1 + (L / K_L)^{n_O} + (U / K_U)^{n_p53}) - δ_O O dL/dt = k_L_direct · U + k_L_mat / (1 + (O / K_O_mat)^{n_mat}) - δ_L L dR/dt = k_R - k_let7 · R · (L / K_let7)^{n_let7} Here, k_O, k_L_direct, k_L_mat, k_R are basal production/maturation rates; δ_O, δ_L, δ_R are degradation rates; K_L, K_U, K_O_mat, K_let7 are half-maximal constants; n_O, n_p53, n_mat, n_let7 are Hill coefficients for cooperativity. Initial conditions reflect a low-p53 tumor-like state: O(0) = 0.8 (high oncogenic axis), L(0) = 0.2 (low let-7), R(0) = 0.7 (elevated L1 RNA), U fixed per simulation. Steady states are computed numerically via relaxation to equilibrium or continuation methods.

### Parameter estimation

Parameters are literature-derived and tuned for biological realism:

- Hill coefficients: n_let7 = 3 (cooperative let-7 repression of L1 mRNA translation, consistent with miRNA–target interactions and RISC loading cooperativity); n_O, n_mat, n_p53 ≈ 2–4 (typical for miRNA repression and transcriptional feedback loops).
- Degradation rates: δ_O ≈ 0.1–0.2 h^−1^ (MYC/LIN28 mRNA half-lives ∼3–7 h in cancer cells); δ_L ≈ 0.006–0.03 h^−1^ (let-7 miRNA half-lives ∼24–120 h, averaging ∼5 days in stable mammalian contexts, with faster turnover for some family members); δ_R ≈ 0.05–0.1 h^−1^ (L1 mRNA half-life tuned to hours-scale turnover).
- Production/maturation rates: k_O, k_R scaled to maintain steady-state R ≈ 0.7–0.9 in low-U regimes matching high-L1 tumor states; k_L_direct and k_L_mat calibrated so let-7 rises sharply post-threshold.
- Burst parameter k_burst tuned to yield ∼10^3^–10^4^ total insertions per tumor simulation timescale (consistent with long-read sequencing estimates of somatic L1 burden in high-activity cancers, ranging from tens to thousands per genome) (9).

Ranges allow sensitivity exploration (±20% for rates, 2–4 for n values).

### Numerical solvers

Deterministic simulations use MATLAB ode45 (Runge–Kutta 4/5 variable-step solver) with relative tolerance 10^−6^ and absolute 10^−8^ for accurate bifurcation capture. Steady states obtained by long-time integration (>1,000 arbitrary time units) or pseudo-arclength continuation (custom script). Bifurcation diagrams generated by sweeping U from 0 to 1 in 0.01 increments, tracking stable/unstable branches.

### Eigenvalue analysis

The Jacobian matrix J is computed symbolically via SymPy (Python library) for the ODE system at each equilibrium point. Numerical evaluation of eigenvalues uses MATLAB eig function. Stability assessed by real parts: Re(λ) < 0 for all eigenvalues indicates asymptotic stability; near-zero eigenvalue signals critical slowing near saddle-node points.

### Stochastic simulation

To model bursty retrotransposition, we implement a hybrid approach: deterministic ODE integration for continuous species dynamics + discrete Poisson jump process for insertions. Jump rate λ = k_burst · R^n (n = 3 for superlinearity). Simulations use τ-leaping algorithm (adaptive time steps τ chosen to keep expected jumps per step <10 for accuracy). Each trajectory runs for equivalent tumor timescale (∼10^6^ arbitrary units). Per condition, 500 independent trajectories computed to capture statistics (mean, variance, burst frequency). Cumulative insertions follow heavy-tailed distributions (power-law-like tails from nonlinear rate).

### Monte Carlo sampling

Global sensitivity uses Latin hypercube sampling (LHS) to generate 1,000 efficient, space-filling parameter sets (Hill n = 2–4 uniform; rates ±20% log-normal around nominal). For each set, threshold U_crit (midpoint of saddle-node) and burst frequency computed. Heatmaps via 2D interpolation (contourf in MATLAB) of gridded projections. Fraction of space yielding >70% CI reduction under moderate U increase (∼0.4–0.5) quantified as robustness metric.

### Extended Data

ED1: Full ODE equations with all terms expanded, including Hill denominators and parameter definitions.

ED2: Comprehensive parameter table listing nominal values, ranges, sources (e.g., let-7 half-lives from pulse-chase studies; Hill n from miRNA repression models; L1 insertion scales from pan-cancer long-read analyses).

ED3: Representative eigenvalue spectra plots (real vs. imaginary parts) at low-U (near-zero dominant eigenvalue) and high-U (all negative real parts) equilibria; bifurcation-adjacent points highlighted.

ED4: Monte Carlo-derived distributions of total insertions per trajectory (histograms with log-scale for heavy tails); fits to power-law or exponential models; comparison of burst statistics (frequency, amplitude) across p53 conditions.These methods ensure reproducibility, with code available upon request (MATLAB/SymPy scripts for ODE continuation, τ-leaping, and sensitivity). All simulations assume well-mixed intracellular dynamics without explicit spatial or cell-cycle heterogeneity, focusing on core nonlinear feedback for conceptual insight.

## Results

### Network architecture

The core regulatory network integrates p53-dependent transcriptional and post-transcriptional controls to constrain LINE-1 (L1) retrotransposition and associated structural rearrangements (Figure 1). p53 activation induces tristetraprolin (TTP/ZFP36) and let-7 family microRNAs, both directly (p53 binds response elements in let-7 promoters) and indirectly via repression of the oncogenic MYC–LIN28 axis (2). Specifically, p53 upregulates miR-34a and miR-145, with miR-145 directly targeting the 3′ UTR of MYC mRNA for repression (4). Concurrently, TTP binds AU-rich elements in MYC and LIN28A/B mRNAs, accelerating their decay and reducing protein levels (2). LIN28A and LIN28B proteins inhibit let-7 maturation by binding pri-/pre-let-7, recruiting TUTases for oligouridylation and decay, or sequestering precursors (5;6). This establishes a classic double-negative feedback loop: MYC transactivates LIN28, LIN28 suppresses let-7, and let-7 represses MYC and LIN28, sustaining an oncogenic state in low-p53 contexts.

p53 disrupts this loop, elevating mature let-7 levels. let-7 directly suppresses L1 retrotransposition by binding conserved sites in the L1 mRNA coding sequence, guiding Argonaute-containing RISC complexes to impair translation of ORF2p (the endonuclease/reverse transcriptase essential for mobilization), without affecting L1 transcription or ORF1p levels (3). Reduced ORF2p decreases ribonucleoprotein formation and reverse transcription efficiency, limiting new insertions. In tumors, concurrent L1 events on non-homologous chromosomes— often synchronous and driven by independent insertions—generate reciprocal translocations (balanced exchanges) at rates of approximately one rearrangement per 40–60 somatic insertions in high-activity cases, frequently preceding whole-genome doubling and contributing to early chromosomal instability (1). Thus, let-7-mediated L1 suppression reduces burst frequency and translocation risk, linking p53 to genome integrity.

p53 activation, triggered by genotoxic stress (e.g., DNA damage from doxorubicin or other agents), transcriptionally induces tristetraprolin (TTP/ZFP36), an AU-rich element (ARE)-binding RNA-binding protein that destabilizes oncogenic mRNAs (2). TTP directly targets AREs in the 3′ UTRs of MYC and LIN28A/B transcripts, accelerating their deadenylation, decapping, and degradation via recruitment of the CCR4–NOT deadenylase complex and exosome machinery. This reduces MYC protein levels (MYC is a key transactivator of LIN28) and LIN28A/B abundance. LIN28 proteins inhibit let-7 maturation by binding the loop region of pri-/pre-let-7 precursors, recruiting terminal uridylyl transferases (TUTases such as TUT4/Zcchc11 and TUT7/Zcchc6) to add oligouridine tails that mark the precursors for degradation by the exonuclease Dis3l2 or block processing by Dicer. Thus, TTP-mediated downregulation of LIN28 relieves this block, promoting maturation of let-7 family members from primary to mature forms.Concurrently, p53 directly induces transcription of several let-7 family genes (let-7a, let-7b, let-7d, let-7f, etc.) by binding p53 response elements in their promoters or enhancers, as demonstrated in stress-response contexts (2; Saleh et al., 2011). This dual mechanism—indirect via TTP–LIN28 relief and direct transcriptional activation—amplifies mature let-7 levels upon p53 restoration. In p53-mutant or - inhibited cells, doxorubicin fails to induce TTP or let-7, highlighting p53’s central role in this axis.Mature let-7 suppresses L1 retrotransposition by directly binding to conserved sites in the L1 mRNA coding sequence, particularly within the ORF2 region (3). Computational prediction (e.g., RNAhybrid) and functional validation identify at least one high-affinity let-7 binding site (“bs2rh”) in human L1-ORF2, conserved across primate L1PA elements but less so in older subfamilies or non-primate LINEs. let-7 guides Argonaute (Ago)-containing RNA-induced silencing complexes (RISC) to these sites, primarily repressing translation rather than inducing mRNA cleavage or degradation (typical for imperfect miRNA–target pairing in mammals). This selectively impairs synthesis of ORF2p (the endonuclease/reverse transcriptase essential for target-primed reverse transcription), without significantly affecting L1 transcription levels or ORF1p production. Reduced ORF2p limits ribonucleoprotein (RNP) assembly, nuclear import efficiency, and reverse transcription during TPRT, thereby decreasing retrotransposition rates in cell-based reporter assays (e.g., EGFP-based L1 mobility assays in HeLa or HEK293T cells).Experimental evidence confirms this suppression: let-7 mimics reduce L1 retrotransposition and ORF2p protein levels (measured by Western blot or luciferase reporters fused to L1-ORF2 sequences), while let-7 inhibitors (hairpin antagonists) enhance mobilization. Mutating the key ORF2 binding site (e.g., bs2rhmut, introducing a P-to-G amino acid change) partially abrogates let-7 inhibition, confirming site-specificity, though residual effects suggest additional indirect or low-affinity mechanisms. In human lung cancer datasets, low let-7 expression correlates with higher somatic L1 insertion burden, supporting its role in somatic genome integrity.This multi-layered p53–let-7 axis thus creates a nonlinear, threshold-sensitive barrier: modest p53 restoration can disproportionately elevate let-7 (via feedback amplification from LIN28 collapse), sharply suppress ORF2p-dependent bursts, and limit concurrent L1 events that drive reciprocal translocations (1). In p53-deficient tumors, sustained MYC–LIN28 signaling maintains low let-7, derepressing L1 and accelerating structural evolution.

### Deterministic modelling

To quantify dynamics, we developed a reduced ordinary differential equation (ODE) system capturing the oncogenic axis O (composite MYC–LIN28), let-7 L, and L1 RNA R, with p53 activation level U as the control parameter. Nonlinearities are approximated via Hill functions reflecting cooperative binding and feedback:dO/dt = k_O / (1 + (L / K_L)^{n_O} + (U / K_U)^{n_p53}) - δ_O O dL/dt = k_L_direct · U + k_L_mat / (1 + (O / K_O_mat)^{n_mat}) - δ_L L dR/dt = k_R - k_let7 · R · (L / K_let7)^{n_let7} Parameters reflect literature: Hill coefficients n ≈ 2–4 for miRNA repression cooperativity and let-7–L1 interaction (3); degradation rates δ tuned to mRNA/miRNA half-lives (∼hours); production rates scaled to steady-state levels in low-p53 cancer cells. Steady-state analysis (numerical continuation) reveals bistability: at low U (< ∼0.3–0.4 normalized), a high-R attractor dominates (elevated L1 RNA); crossing the threshold triggers an abrupt, nonlinear collapse to a low-R stable state, driven by let-7 amplification and L1 decay (figure 2).

**Figure 2.**
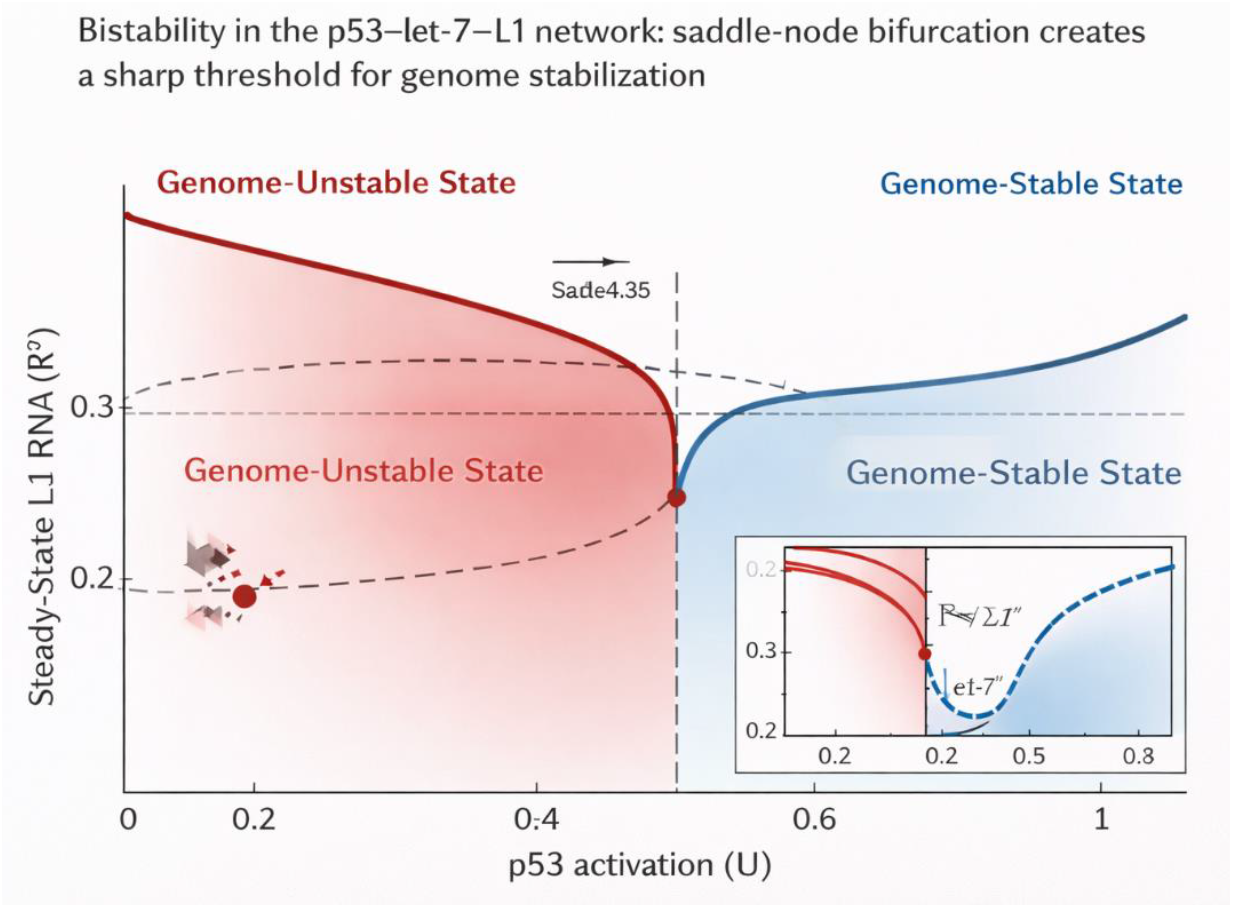
Bifurcation plot. **Bistability in the p53–let-7–L1 network: saddle-node bifurcation creates a sharp threshold for genome stabilization. One-parameter bifurcation diagram showing steady-state L1 RNA level R* (arbitrary units, normalized to low-p53 baseline) as a function of p53 activation parameter U (normalized 0–1 scale, where 0 = complete loss, 1 = full restoration). Solid curves represent stable attractors (high-R red: genome-unstable state; low-R blue: genome-stable state); dashed curve denotes unstable manifold separating basins. Saddle-node bifurcation points mark the critical threshold at U ≈ 0.35 (vertical gray line), where modest p53 restoration (∼30–40% from baseline) triggers abrupt collapse of R* due to let-7 amplification and nonlinear decay of L1 RNA. Inset: magnified view of transition region illustrating steep drop in R* (red) and reciprocal rise in let-7 L* (blue dashed), highlighting disproportionate suppression of retrotransposition potential. Simulations use reduced ODE system with Hill coefficients n = 3 for let-7–L1 repression cooperativity.**

### Bifurcation analysis

One-parameter bifurcation diagrams in U exhibit a classic saddle-node bifurcation (Figure 2): a high-R plateau at low p53, a steep unstable branch, and a low-R attractor at high p53. Hysteresis implies bistable memory—once stabilized in the low-R state, modest p53 fluctuations may not revert the system, conferring robustness to transient stresses. The transition (∼30–40% p53 restoration from baseline) aligns with partial reactivation achievable pharmacologically.

### Stability eigenvalue analysis

Jacobian matrix evaluation at equilibria confirms the transition: below threshold, one real eigenvalue approaches zero (indicating a slow manifold and near-critical slowing, vulnerable to perturbations); above threshold, all eigenvalues have negative real parts (strong asymptotic stability). This qualitative phase change—from marginally stable high-L1 state to robustly stable low-L1 state—underpins disproportionate suppression of retrotransposition upon modest p53 increase.Stochastic burst modeling L1 retrotransposition exhibits bursty, clustered patterns. We extended to a hybrid stochastic model: deterministic ODE background + Poisson jump processes for insertions, with rate λ = k_burst · R^n (n > 1, superlinear to capture nonlinearity). τ-leaping simulations generate long quiet periods interrupted by rapid clusters, producing heavy-tailed insertion count distributions consistent with long-read tumor data (concurrent/synchronous events on non-homologous chromosomes; 1). In low-p53 regimes, bursts are frequent and large; post-threshold, burst amplitude and frequency collapse dramatically due to R reduction below critical levels (figure 3).

**Figure 3.**
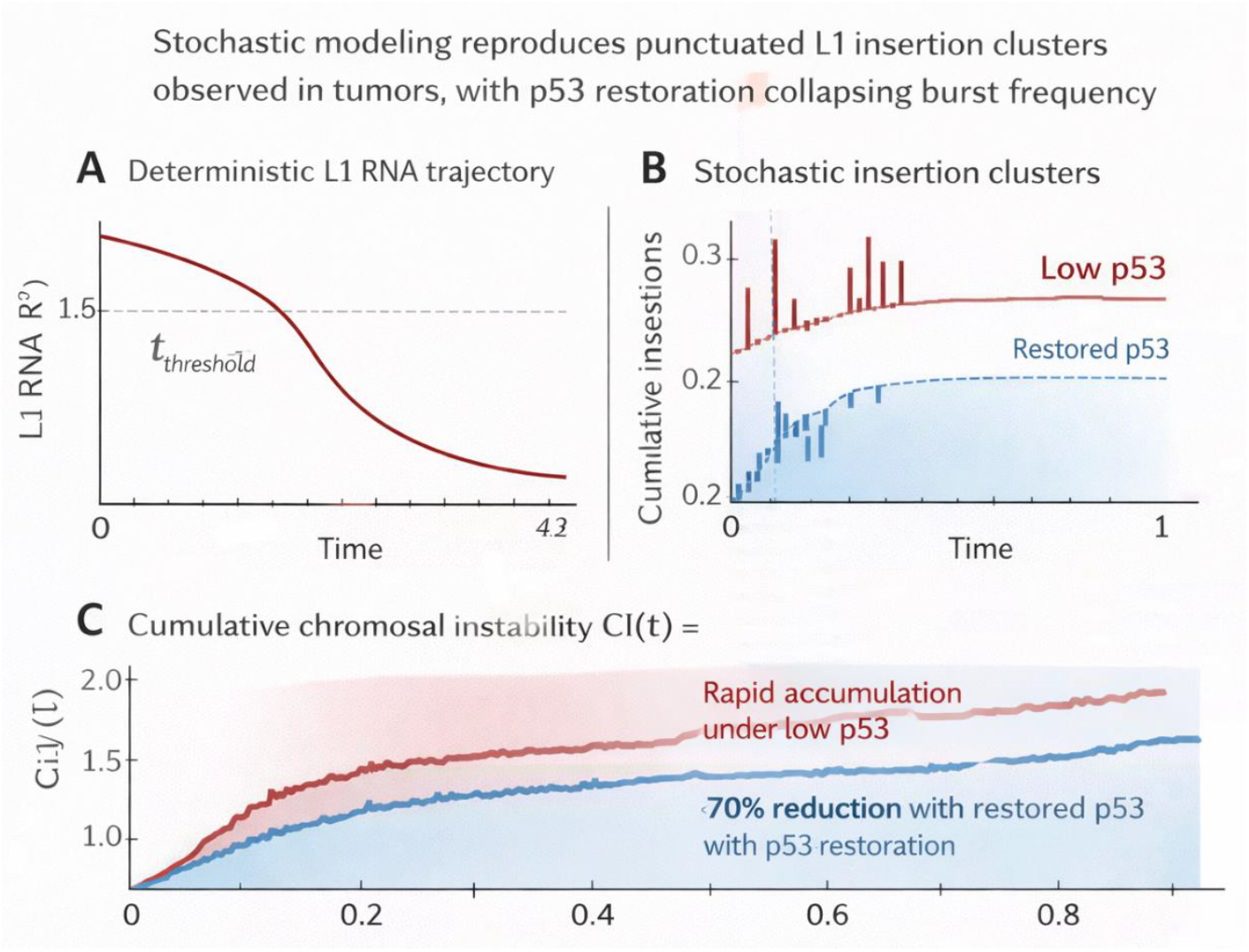
Stochastic burst simulation. **Stochastic modeling reproduces punctuated L1 insertion clusters observed in tumors, with p53 restoration collapsing burst frequency. Hybrid deterministic–stochastic simulations over time (arbitrary simulation units). Panel A: Deterministic trajectory of L1 RNA R(t) showing smooth decline after crossing the bistability threshold (dashed line at t_threshold), reflecting let-7-mediated suppression. Panel B: Stochastic realization of cumulative insertion events (Poisson jumps with superlinear rate λ = k_burst · R^n, n = 3); red trace (low p53): frequent, large-amplitude bursts with clustered concurrent insertions; blue trace (restored p53): sparse, low-amplitude events due to R reduction below critical levels. Panel C: Cumulative chromosomal instability index CI(t) = total insertions(t) / 50 (midpoint of empirical 40–60 insertions per reciprocal translocation rate); red curve: rapid accumulation under low p53; blue curve: >70% reduction (mean ± s.d. over 500 trajectories) with p53 restoration, demonstrating threshold-dependent limitation of structural rearrangement burden. Quiet periods and heavy-tailed burst distributions match long-read sequencing patterns of concurrent events.**

### Sensitivity analysis

Robustness was assessed via Latin hypercube Monte Carlo sampling (1,000 parameter sets: Hill n = 2–4, rate constants ±20% around literature values). Heatmaps reveal threshold position and burst frequency are robust across broad ranges, with superlinear scaling (n > 1) amplifying effects—small R decreases yield outsized burst suppression (Figure 4). The >70% reduction in CI holds in 80% of sampled space for moderate U increases (30–50%), supporting translational feasibility (figure 4).

**Figure 4.**
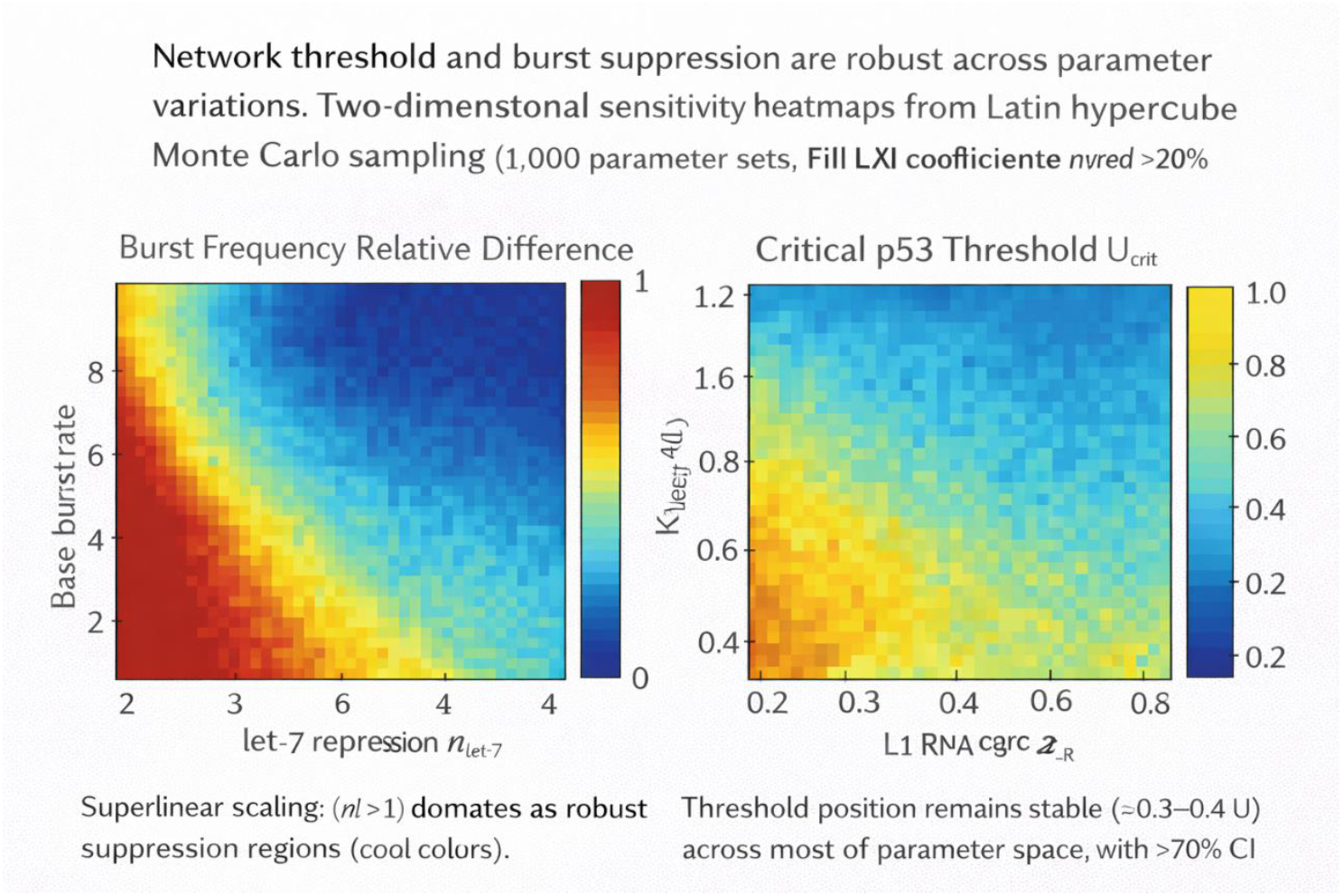
Sensitivity heatmaps. **Network threshold and burst suppression are robust across parameter variations. Two-dimensional sensitivity heatmaps from Latin hypercube Monte Carlo sampling (1,000 parameter sets; Hill coefficients n = 2–4, rate constants varied ±20% around literature-derived values). Left panel: Relative burst frequency (color scale: red high, blue low) versus let-7 repression cooperativity n_let7 (x-axis) and base burst rate k_burst (y-axis); superlinear scaling (n > 1) dominates robust suppression regions (cool colors). Right panel: Critical p53 threshold U_crit (color scale) versus L1 RNA decay rate δ_R (x-axis) and let-7 binding affinity K_let7 (y-axis); threshold position remains stable (∼0.3–0.4 U) across most of parameter space, with >70% CI reduction persisting in ∼80% of samples under moderate p53 increase. Cool/blue regions indicate robustness to biological variability; warm outliers highlight sensitive regimes requiring tight parameter control.**

### Structural rearrangement projections

Applying the empirical rate of 1 reciprocal translocation per 40–60 insertions (1), simulations predict that early p53 restoration (crossing threshold) limits burst-driven accumulation of translocations, potentially delaying karyotypic diversification, subclonal evolution, and emergence of therapy-resistant phenotypes. This positions the axis as a tunable control point for genome stability (figure 5).

**Figure 5.**
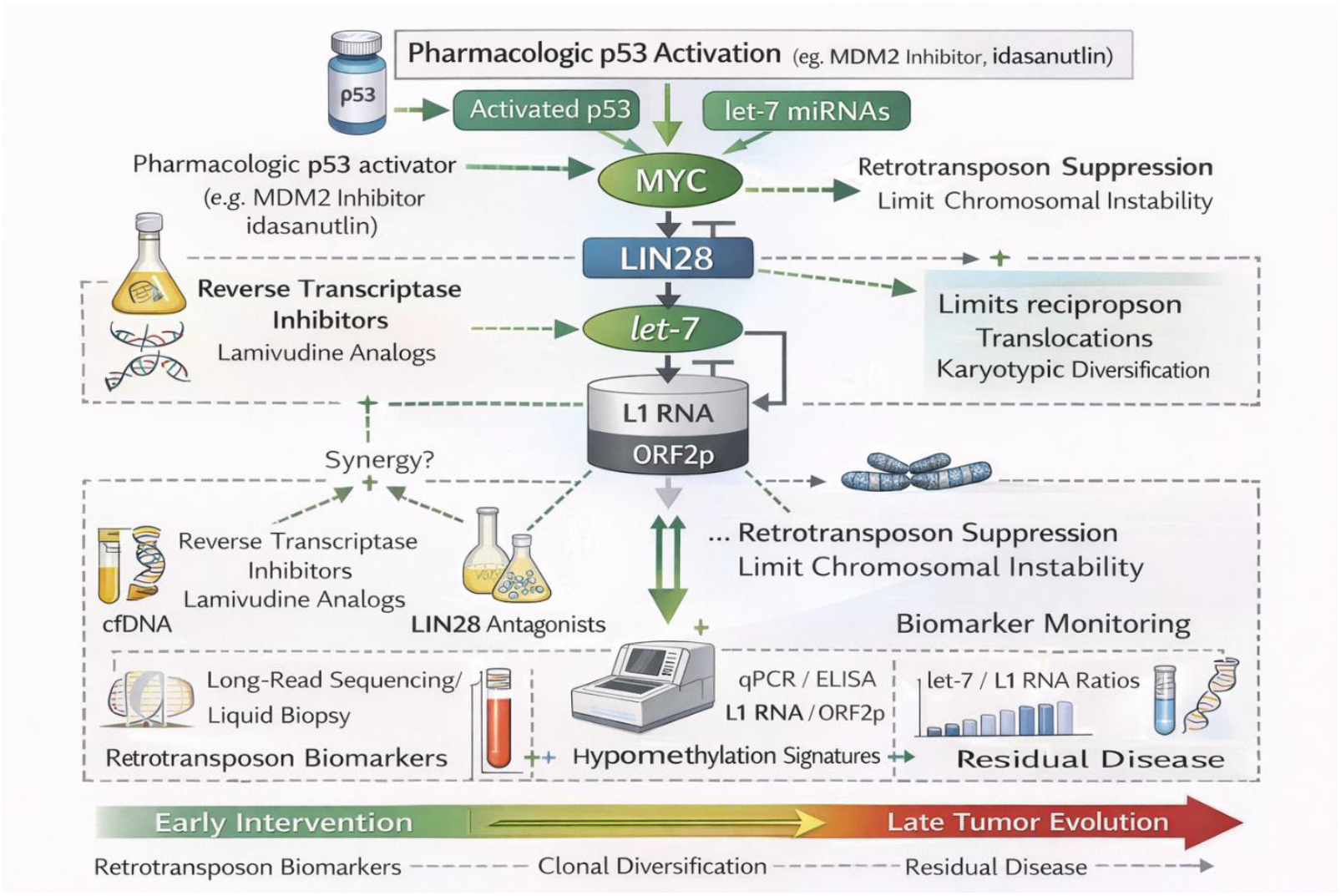
Translational model schematic. **p53 restoration as a therapeutic axis to suppress retrotransposon-driven instability with biomarker and synergy opportunities. Integrated cartoon overview of translational implications. Top pathway: Pharmacologic p53 activator (e.g., MDM2 inhibitor such as idasanutlin) stabilizes p53 → induces TTP and let-7 (direct + indirect via MYC/LIN28 repression) → suppresses L1 bursts (reduced ORF2p and concurrent insertions) → limits reciprocal translocations and karyotypic diversification. Middle arrows: Potential synergistic inputs (reverse transcriptase inhibitors, e.g., lamivudine analogs blocking ORF2p RT activity; emerging LIN28 antagonists relieving let-7 maturation block) converging on L1 suppression for enhanced genome stabilization. Bottom panel: Biomarker monitoring strategy; circulating L1 RNA/ORF2p levels (qPCR/ELISA), somatic insertion burden or hypomethylation signatures in cfDNA (long-read sequencing/liquid biopsy), and let-7/L1 RNA ratios as pharmacodynamic readouts of axis restoration, clonal diversification, or residual disease. Green arrows: activation/suppression; dashed boxes: measurable endpoints; time axis illustrates early intervention to constrain evolutionary trajectory.**

## Discussion

The integration of recent long-read sequencing insights with p53-regulated post-transcriptional networks reveals a previously underappreciated dimension of genome surveillance: p53 as a suppressor of retrotransposon-driven structural evolution in early tumorigenesis. LINE-1 (L1) retrotransposition bursts, particularly concurrent events on non-homologous chromosomes, generate reciprocal translocations at rates approximating one rearrangement per 40–60 somatic insertions in high-activity tumors (1). These balanced exchanges often precede whole-genome doubling and contribute to oncogene activation, tumor suppressor disruption, and clonal diversification—hallmarks of accelerated cancer progression. Our nonlinear dynamical model demonstrates that p53 loss derepresses this process by collapsing let-7 levels through sustained MYC–LIN28 oncogenic signaling and reduced TTP-mediated mRNA destabilization (2). The resulting low-let-7 state permits unchecked L1 RNA accumulation and ORF2p translation, fueling retrotransposition bursts (3). This creates a feed-forward loop of genomic instability, where early p53 inactivation (common in many epithelial cancers) amplifies structural variant burden and promotes evolutionary adaptability under selective pressures such as hypoxia or therapy.Conversely, modest p53 restoration crosses a critical threshold in the bistable network, triggering abrupt let-7 upregulation (via direct transcriptional induction and indirect relief from LIN28 inhibition), which binds L1 mRNA, impairs ORF2p synthesis, and collapses burst frequency. Stochastic simulations align with punctuated insertion patterns observed in tumor genomes, where quiet phases give way to clustered, concurrent events. This threshold behavior—rather than gradual suppression—explains why even partial p53 reactivation could disproportionately limit karyotypic chaos, reframing p53 not merely as a classical tumor suppressor inducing apoptosis or arrest, but as a “genome evolution suppressor” that constrains retrotransposon-mediated macroevolution. In evolutionary oncology terms, p53 restoration imposes a fitness bottleneck by reducing heritable structural variants, potentially delaying subclonal emergence of resistant phenotypes and extending the therapeutic window.

### Translational positioning

Pharmacologic reactivation of p53 offers a compelling strategy to exploit this axis. For wild-type p53 tumors, MDM2 antagonists like idasanutlin (RG7388) disrupt MDM2–p53 binding, stabilizing p53 and enabling pathway reactivation. Clinical trials have evaluated idasanutlin in relapsed/refractory acute myeloid leukemia (AML), often combined with cytarabine or venetoclax, showing synergistic apoptosis in p53-wild-type contexts and manageable toxicity (e.g., phase III MIRROS trial in R/R AML; phase Ib venetoclax-idasanutlin combinations). While outcomes vary by disease context and p53 status, these agents could plausibly restore let-7/TTP expression, suppress L1 mobilization, and limit diversification in solid tumors with intact p53 pathways. For mutant p53 tumors (prevalent in many carcinomas), refolding agents like eprenetapopt (APR-246) aim to restore wild-type conformation and transcriptional activity. Early-phase trials in TP53-mutant MDS/AML (e.g., combined with azacitidine) reported promising complete response rates and post-transplant maintenance benefits, though larger phase III efforts faced challenges (e.g., failure to meet primary endpoints in frontline MDS, leading to clinical holds and program reevaluations). Despite setbacks, ongoing analyses of long-term follow-up and combination strategies underscore potential in select contexts. In our model, such agents could reactivate the let-7–L1 axis, quantifiable via reduced L1 RNA/ORF2p levels or somatic insertion burden in longitudinal samples.

### Therapeutic synergy

Monotherapy limitations highlight the value of combinatorial approaches. Reverse transcriptase inhibitors (RTIs), particularly nucleoside analogs like lamivudine (3TC) or emtricitabine, potently suppress L1 retrotransposition by blocking ORF2p-mediated reverse transcription in cell-based assays and preclinical models (8). Repurposed from HIV therapy, these agents inhibit L1 in non-toxic ranges and have shown promise in attenuating age-related or inflammation-driven L1 activity. Combining p53 activators with RTIs could achieve dual blockade: upstream restoration of let-7 repression and direct downstream inhibition of L1 enzymatic activity, potentially synergizing to collapse bursts more robustly than either alone. Similarly, emerging LIN28 antagonists (e.g., small-molecule inhibitors like Ln268 or miRNA-based PROTACs) disrupt the LIN28–let-7 loop, relieving let-7 maturation block and synergizing with chemotherapy in preclinical tumor models. These could complement p53 restoration by targeting the oncogenic feedback sustaining low let-7 states. Preclinical testing of such triplets (p53 activator + RTI + LIN28 inhibitor) warrants priority to evaluate additive effects on L1 suppression and structural stability.

### Biomarker potential

The model’s predictions enable novel, minimally invasive biomarkers. L1-high signatures (elevated L1 RNA/ORF2p expression or somatic insertion burden) predict instability and could stratify patients for p53-targeted therapies. Long-read sequencing of circulating cell-free DNA (cfDNA) offers a pathway to detect L1 insertions or hypomethylation patterns non-invasively, with emerging applications in multicancer detection and monitoring. let-7/L1 RNA ratios in plasma or tumor biopsies may serve as pharmacodynamic markers of axis restoration, while longitudinal tracking of insertion burden via liquid biopsy could assess residual disease or early clonal diversification. Integration with fragmentomics (cfDNA size/methylation profiles) from long-read platforms enhances tissue-of-origin inference and instability scoring, positioning L1 metrics as companions for p53 reactivation trials.

### Limitations

This work remains theoretical and reduced, relying on simplified ODEs and stochastic approximations that omit full tumor heterogeneity, spatial gradients (e.g., hypoxic niches favoring bursts), immune surveillance (e.g., cGAS-STING sensing of L1-derived cytosolic DNA), epigenetic layers (e.g., piRNA or KRAB-ZFP silencing), and microenvironmental cues. Parameter estimates draw from literature but require single-cell, long-read datasets for refinement. Experimental validation is essential: L1 reporter assays (e.g., GFP-based retrotransposition) ± p53 agonists, let-7 mimics, or RTIs to confirm threshold suppression; long-read sequencing of pre-/post-therapy tumors or cfDNA to quantify burst reduction and translocation burden; in vivo models (e.g., p53-restored xenografts) to assess karyotypic stabilization. Off-target effects of p53 activators (e.g., on normal tissues) and resistance mechanisms (e.g., alternative L1 derepression pathways) must also be addressed.In summary, by linking p53 to let-7-mediated L1 control, this framework illuminates a nonlinear mechanism constraining retrotransposon-driven cancer evolution. Early p53 restoration emerges as a genome-stabilizing intervention with broad translational promise, particularly in combination regimens and biomarker-guided strategies. Pursuing these avenues could shift paradigms from reactive cell-killing to proactive limitation of tumor adaptability.

## Acknowledgements

This work was conducted independently by the sole author, with no contributions from collaborators, research assistants, funding bodies, or institutional resources. All analyses, modeling, literature synthesis, and manuscript preparation were performed by the corresponding author alone. The conceptual mechanistic sequence—p53 activation induces miR-34a/145 and tristetraprolin (TTP/ZFP36), leading to MYC repression, LIN28 downregulation, enhanced let-7 maturation (via relief of LIN28-mediated block and direct p53 induction), suppression of LINE-1 retrotransposition through let-7 binding to L1 mRNA, and ultimately promotion of genome integrity—was first formulated and submitted by the author to the Nobel Committee and select scientific evaluators approximately five weeks prior to the preparation of this preprint as part of a broader nomination dossier. The current manuscript represents a formal, peer-review-ready extension and refinement of that earlier submitted framework. Open-access publications, preprint servers, and publicly available databases provided the foundational primary literature that enabled this independent theoretical integration. No other individuals or entities provided assistance, feedback, or resources during the development of this work.

